# Exploring Neuroscience Researchers’ Trust in Preprints through Citation Analysis

**DOI:** 10.1101/2024.04.29.591455

**Authors:** Behrooz Rasuli, Fatemeh Seyfzadehdarabad, Paolo Cardone, Aurore Thibaut, Olivia Gosseries

## Abstract

Preprints have emerged as efficient tools for fast and free dissemination of scientific findings. The present study explores the evolving landscape of preprints among the field of neuroscience and examines patterns and evolution of citations to preprints over time in this field. Leveraging bibliometric methods, we identified over 33,000 citations (1993-2022) to preprints within neuroscience publications indexed in Scopus. The findings elucidate a significant temporal increase in the number of documents citing preprints, reaching a peak of around 60 per 1,000 Scopus documents in 2021. Diverse document types, particularly reviews, exhibit a growing reliance on preprints as references. The most frequently cited preprint servers include bioRxiv, ArXiv, medRxiv, and PsyArXiv. Leading journals such as eLife and PLOS Computational Biology have cited preprints more than others. The United States takes the lead in citing preprints, followed by the United Kingdom and Germany. Using Scite.ai, motivations underlying preprint citations and the context in which they were cited were explored and the results are indicative of the “mentioning” nature of 93% of citations to preprints. Furthermore, the introduction and discussion sections have shown to include the highest number of citations to preprints. The findings highlight the dynamic transformation of preprints in neuroscience field.

## 1 Introduction

### 1.1 Background

A preprint is defined as “*a complete version of an original manuscript posted by the authors to an open-access server before formal peer-review process*” (Alfonso & Crea, 2023). Preprinting is considered as an efficient and successful means of timely and free of charge dissemination of scientific findings (Sever et al., 2019). In line with open access principles of open and free dissemination of research outputs, preprint servers provide the public good to “everyone”, which can be depicted against heavy open access publication charges. In fact, the open access movement was initiated as a consequence of the “serials crisis” (Young, 2009), which posed challenges for the academic libraries with limited budgets. The problem, however, persists with researchers now carrying the financial burdens. This becomes especially pronounced with policies adopted by funding organizations, universities, and health-related institutes, requiring open access publication of the research data and findings for a wider accessibility and a higher visibility (Voronin et al., 2011).

With preprinting, financial resources can be better managed and allocated, and open and easy access to research findings will be promoted. Nevertheless, not only are there still objections towards it, not many researchers are even aware of such a scholarly communication (Rong & Ludo, 2023). On an apparent negative note, preprints do not go through the traditional formal peer-review and this is one of the main reasons behind trust issues towards preprints (Arnold et al., 2021; Rong & Ludo, 2023). In relation to this, an analysis of citations to preprints in PLOS journals has shown that preprints are more frequently cited in the “methods” section as compared to other sections of an article (Bertin & Atanassova, 2022); however, different lexical contents (i.e., terminology and linguistic characteristics) are used for citations to emphasize that they are so-called “non-validated” preprints and not peer-reviewed publications (Bertin & Atanassova, 2022). Although peer-review is considered as a standard quality control process, peer-reviewed publication can also be subject to several drawbacks, including inordinate wait times and reliance on (potentially biased) human judgment (Wingen et al., 2022). Along the same lines, lack of quality control on preprint servers can be considered as a misconception since most of the preprint servers have quality control systems as illustrated by four to five days turnaround time on medRxiv for a more careful scrutiny (Kwon, 2020). Although the vetting systems on preprint servers are far more basic than those of publication venues with peer-review, this flaw cannot be considered as a *hamartia* (referring to Aristotle’s concept of the “fatal flaw” leading to the downfall of the tragic hero/heroine [Devi, 2014]) as with preprints, the reader — not the referees — will decide if a piece of research is worth being studied and cited.

In the same context, to address researchers’ skepticism towards preprints due to lack of review and to avoid serious consequences of health-related findings, “bioRxiv” and “medRxiv” preprint servers improved their screening procedures through blocking speculative research works during the Covid-19 pandemic (Kwon, 2020). The skepticism exists while a study on preprints published during a 30-year period showed that 41% of preprints were subsequently published at a peer-reviewed journal (Xie et al., 2021). Quite interestingly, a recent study comparing preprint and peer-reviewed versions of randomized clinical trials (RCT) on Covid-19 showed that 119 out of 152 (78%) RCT preprints were finally published in a peer-reviewed journal. In 65 out of 119 (55%) peer-reviewed articles, differences were spotted in outcomes, analyses, results, or conclusions, yet the main conclusion remained consistent in 98% of the cases (Bai et al., 2023). Similarly, another study showed the quality of reporting in the biomedical literature is higher in peer-reviewed articles compared to preprints, yet the difference is small (5%) (Carneiro et al., 2020).

Despite a partially biased attitude towards the credibility of preprints, a study on four subject repositories of arXiv, Research Papers in Economics (RePEc), Social Science Research Network (SSRN), and PubMed Central (PMC) through a citation analysis of the Scopus-indexed publications displayed a rapid increase in the number of citations to these subject repositories on an interdisciplinary level (Li et al., 2015). This was further confirmed by another study on arXiv preprints showing preprints in physics and astronomy, mathematics, computer science, and engineering as the most highly cited, compared to other academic areas, including dentistry, veterinary, nursing, psychology, and energy (Noruzi, 2016). On a positive note, preprints contribute to transparency and report non-significant or contradictory findings (da Silva, 2018) without fear of rejection based on (potentially biased) human judgments. On top of that, preprints contribute to higher visibility as previously evidenced that among articles in biomedical journals, those with a preprint (5405 out of 74,239) had a 36% more citations and a 49% higher altmetric attraction score (as reported in (Chaleplioglou & Koulouris, 2021)). In the field of mathematics as well, “early-view” and “open-access” features led to more citations and higher readership (Wang et al., 2020). Interestingly, publication of highly cited physics-related arXiv preprints in a journal is found less likely (Kim et al., 2020).

Although deep concerns hover around the risk of disseminating harmful misinformation and favoring authors over patients (Arnold et al., 2021), preprints can still be instrumental in health care advances depicted by increased use of preprints by biologists and medical communities under emergency conditions (Chaleplioglou & Koulouris, 2021). Given the persistent and sophisticated unknowns of the brain and the huge financial and social burdens associated with increasing neurological disorders (e.g., brain injury, Alzheimer’s) (Centers for Disease Control and Prevention, 2019; GBD 2017 US Neurological Disorders Collaborators, 2021), wider in-depth research activities are required in the emerging and rapidly improving field of neuroscience. For instance, in spite of decades of debates and discussions on the “hard problem of consciousness” (Chalmers, 1995), consciousness still remains a mystery which hinders prognosis and treatment of patients with disorders of consciousness. Therefore, to speed up the process of brain discovery, preprints of neuroscience research findings can, to some extent, contribute to a better understanding of the immense complexity of the brain and potentially a control over brain-related diseases in a shorter period of time.

Neuroscience is an emerging yet rapidly improving field, leading to multifaceted advances in different societal aspects even beyond medicine (Cara et al., 2020), for example in justice where several murder convictions have been reduced to manslaughter with neuroscientific evidence (Catley & Claydon, 2015). To take this into account and to encourage timely, free, and transparent publication of neuroscientific findings at different stages of discovery, investigating the preprinting culture among neuroscience researchers sounds imperative. In this study, thus, we aim to investigate (1) to what extent neuroscience researchers are engaging with preprints and (2) what are their motivations behind citing preprints (supporting, contradicting, or mentioning) to have an impression of the level of trust neuroscientists put in preprints and discuss its implications and applications.

### 1.2 Preprint Servers

Preprints have been a component of the scholarly publishing domain for a long time and the practice of sharing preprints has been in operation since the 1960s (Smart, 2022). However, before the 1990s, with no possibility of instant information sharing without smartphones and social media feeds, physical mails served as the primary method of distributing preprints. In 1991, a physicist named Paul Ginsparg at Los Alamos National Laboratory launched an automated email server specifically for distributing high-energy physics preprints (Berg et al., 2016). This seemingly simple innovation circumvented the physical mail constraints, enabling fast and free dissemination. This email server eventually evolved into arXiv, the inaugural official preprint server and a web service facilitating sharing and accessing preprints globally.

Inspired by arXiv, other specialized preprint servers were developed. In 1994, the Social Science Research Network (SSRN) emerged as a major repository for economics, law, and social sciences (Xie et al., 2021) and in 1997, Research Papers in Economics (RePEc) was launched focusing solely on economics.

Starting from 2010, numerous new preprint servers have been introduced, several of which are supported by the Center for Open Science (COS). This organization initiated the Open Science Foundation (OSF) to serve as a platform facilitating the organization of preprint collections. However, largely increased preprints on Covid-19 in 2020 marked a significant turning point (Smart, 2022). As of December 26-29, 2023, Table 1 provides classified sets of data on the number of records on preprint servers of a wide range of disciplines and regions along with their launch year and maintaining organization, reflecting the dynamic landscape of preprint sharing in the academic community.

**Table 1.**
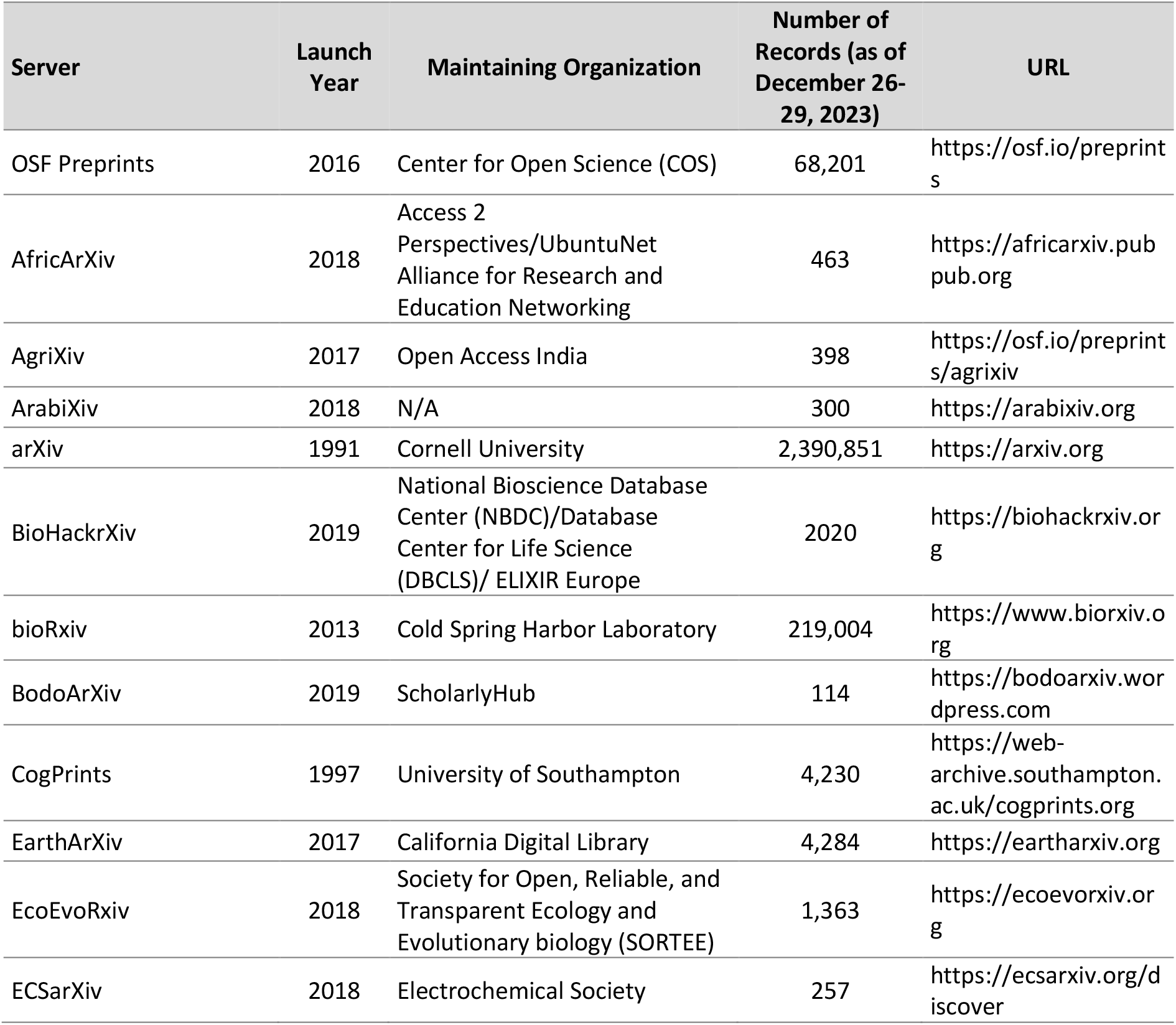
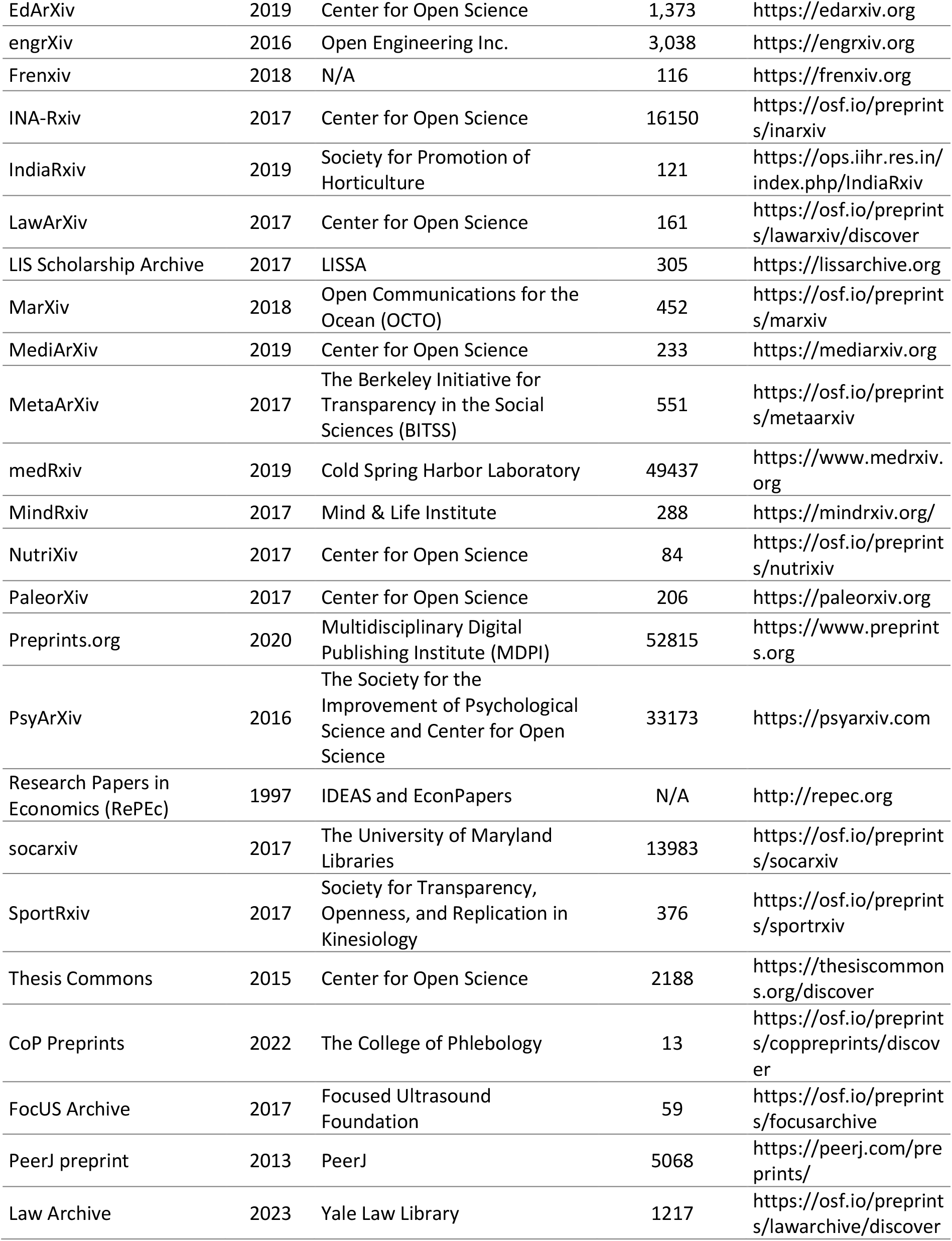
Preprint servers included in the current study.

According to Table 1, a diverse range of organizations, including academic institutions, professional societies, non-governmental organizations, and publishers are maintaining preprint servers. ArXiv is the oldest and largest preprint server with more than 2.3 million records. Certain preprint servers in this table are no longer active. Some preprint servers are hosted by the COS and the OSF Preprints platform, while others have independent hosting servers (The data presented in Table 1 has been compiled from multiple sources, including https://en.wikipedia.org/wiki/List_of_preprint_repositories, https://doapr.coar-repositories.org/repositories, https://www.library.ucsb.edu/research/resources/databases, and the homepages of respective preprint servers).

Preprints, nevertheless, are no longer exclusively published on preprint servers today with numerous institutional repositories emerging as essential hubs for hosting preprints. Yet, preprint servers are still widely recognized and utilized for sharing preprints. To ensure accessibility and discoverability of preprints as well as lasting recognition for these scholarly works, the majority of preprint servers assign a digital object identifier (DOI) to preprints (Moshontz et al., 2021). It should be, however, pointed out that some types of research might not be accepted for publication on certain preprint servers, as is the case with BioRxiv stating “*The bioRxiv subject categories Clinical Trials and Epidemiology are now closed to new submissions*”.

Notably, preprint servers do not engage in peer-review process, yet they follow their own screening procedures before posting a submitted manuscript. These procedures vary across servers, with a basic check conducted by an individual or an artificial intelligence-based tool or a more comprehensive review of the manuscript. For example, according to the medRxiv submission guideline, every manuscript submitted to medRxiv undergoes a preliminary screening process by in-house staff with scientific/editorial expertise and volunteer clinicians and health professionals (known as medRxiv Affiliates) to identify offensive or non-scientific content, as well as material that could potentially pose a health risk (medRxiv, 2019).

## 2 Methods

To address the research questions, the data were collected and analyzed using bibliometric methods and tools. The required data were exported from the Scopus database using a specific query. Scopus was chosen for two main reasons: it has a wide coverage with over 28,000 scholarly source titles and allows users to search within the cited reference fields. The query was designed to retrieve publications in the field of neuroscience containing at least one citation to a preprint (Query 1). The query included the titles of preprint servers listed on the OSF portal (35 server titles, as of September 20, 2023), and medRxiv (an important server hosting preprints related to neuroscience). Additionally, the term “preprint” was appended to certain server titles (i.e., PeerJ, CoP, Law Archive, and FocUS Archive) to avoid potential confusion with journal titles.

### Query 1. The Scopus query for retrieving neuroscience publications with citation(s) to preprint(s)

*REFSRCTITLE (“OSF Preprints” OR “open science foundation Preprints” OR *africarxiv* OR *agrixiv* OR *arabixiv* OR *arxiv* OR *biohackrxiv* OR *biorxiv* OR *bodoarxiv* OR *cogprints* OR *eartharxiv* OR *ecoevorxiv* OR *ecsarxiv* OR *edarxiv* OR *engrxiv* OR *frenxiv* OR “INA-Rxiv” OR *indiarxiv* OR *lawarxiv* OR “LIS Scholarship Archive” OR *marxiv* OR *mediarxiv* OR *metaarxiv* OR mindrxiv OR *nutrixiv* OR paleorxiv OR “Preprints*.*org” OR psyarxiv OR *repec* OR *socarxiv* OR *sportrxiv* OR “Thesis Commons” OR “CoP preprint” OR “FocUS Archive preprint” OR “PeerJ preprint” OR “Law Archive preprint” OR *medrxiv*) AND SUBJAREA (neur) AND PUBYEAR<2023*

The search was conducted on September 20, 2023, and returned a total of 19,941 records which were downloaded as a CSV file and processed. The dataset includes the title, DOI, sources, and EID of these records. The references field of each publication was split using a Python code, resulting in each reference and the EID of the citing document being placed on a separate line in a text file. From this text file, 33,754 references, containing the titles of preprint servers, were extracted using another Python code (Rasuli et al., 2023). If a citation was mentioned more than once in a single document, it was counted multiple times in the analyses. To ensure the relevance of these references, 100 references were randomly checked to avoid references with the title of preprint servers indicated anywhere other than the publication venue. All the 100 references were published on a preprint server. Finally, the name of preprint servers in each reference was identified. Descriptive statistics were used to summarize and describe the collected data.

To gain a deeper understanding of the context and nature of citations to preprints within the text of publications, from the total of 33,754 citations to preprints within neuroscience-related documents, a random sample of 653 citations was selected for further analysis. The sample size was determined using the Survey System tool (https://surveysystem.com/sscalc.htm), with a 99% confidence level and a confidence interval of 5%. For random selection of the sample, the RAND() function was used in MS Excel (allocating a random number to each citation, sorting the random numbers from the largest to the smallest, and extracting the first 653 records as the research sample). The records containing the sampled citations were identified, their DOIs were retrieved, searched on Scite to identify (1) the location within the document where the preprint was cited (e.g., introduction, methods) and (2) the motivation behind citations (mentioning, supporting, or contradicting). Scite web tool was used to objectively categorize citations and identify self-citations. Scite is developed using machine learning and document ingestion methods. It illustrates if the referenced worked is cited in a “supporting”, “contradicting”, or simply “mentioning” manner while considering the surrounding context in the citing paper and using predefined classifications based on a deep learning model. More precisely, “mentioning” citations provide background information, “supporting” citations confirm/support findings of the cited work, and “contradicting” citations challenge the content/findings of the cited work. More than 25 million scientific articles have been analyzed to develop this tool, which has a database of >880 million classified citations now.

To answer our questions more precisely, we provide a more general overview of how preprints are cited in the neuroscience literature by reporting descriptive statistics on how preprints are cited over the past two decades since they first emerged, within different peer-reviewed document types, on preprint servers, in different journals, and across countries.

## 3 Results

Figure 1 illustrates the temporal trend of Scopus-indexed neuroscience publications with at least one citation to a preprint.

**Figure 1.**
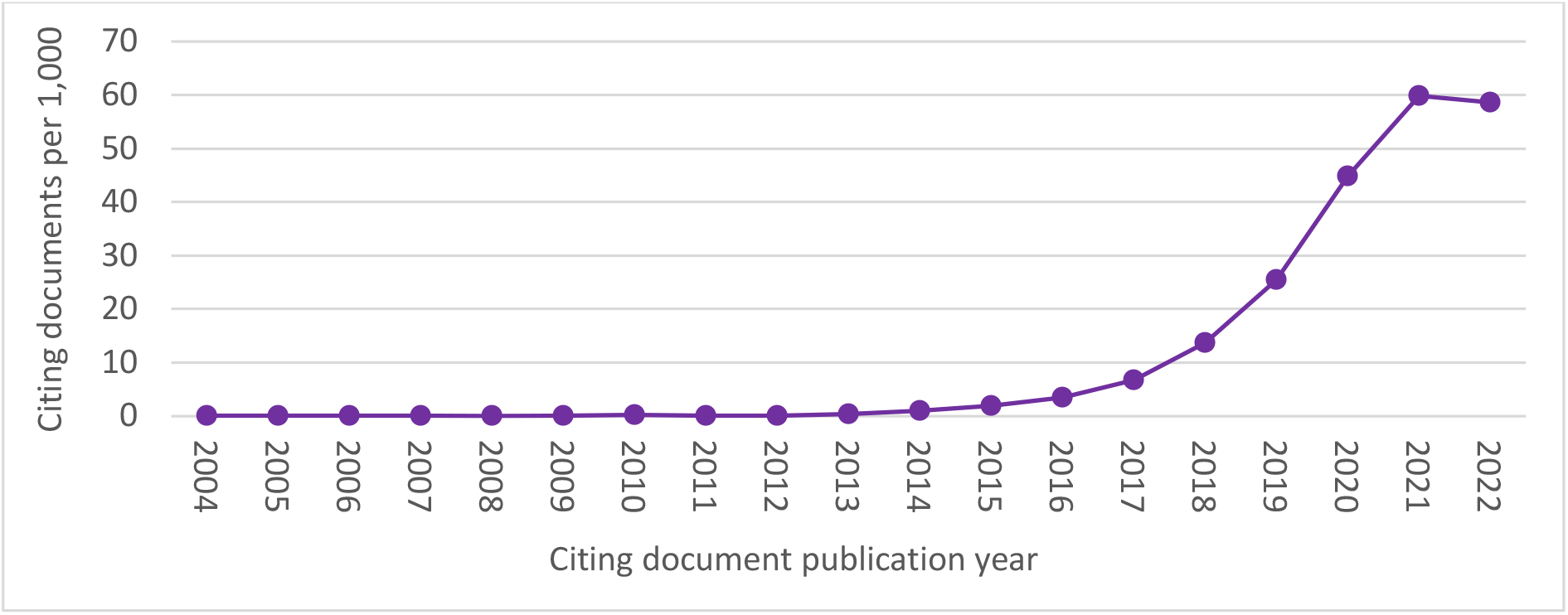
Documents citing preprints per 1,000 Scopus documents.

Figure 1 demonstrates a significant increase in the number of documents citing preprints per 1,000 Scopus documents over time, starting from less than one document per 1,000 documents in the early 2000s. From 2014 onwards, a substantial growth was, however, observed in the number of documents citing preprints. By 2021, the number of citing documents reached a peak of around 60 per 1,000 Scopus documents (6%).

Figure 2 presents an overview of the types of documents citing preprints per 1,000 Scopus documents. The graph categorizes the citing documents into articles, reviews, book and book chapters, notes, and conference papers.

**Figure 2.**
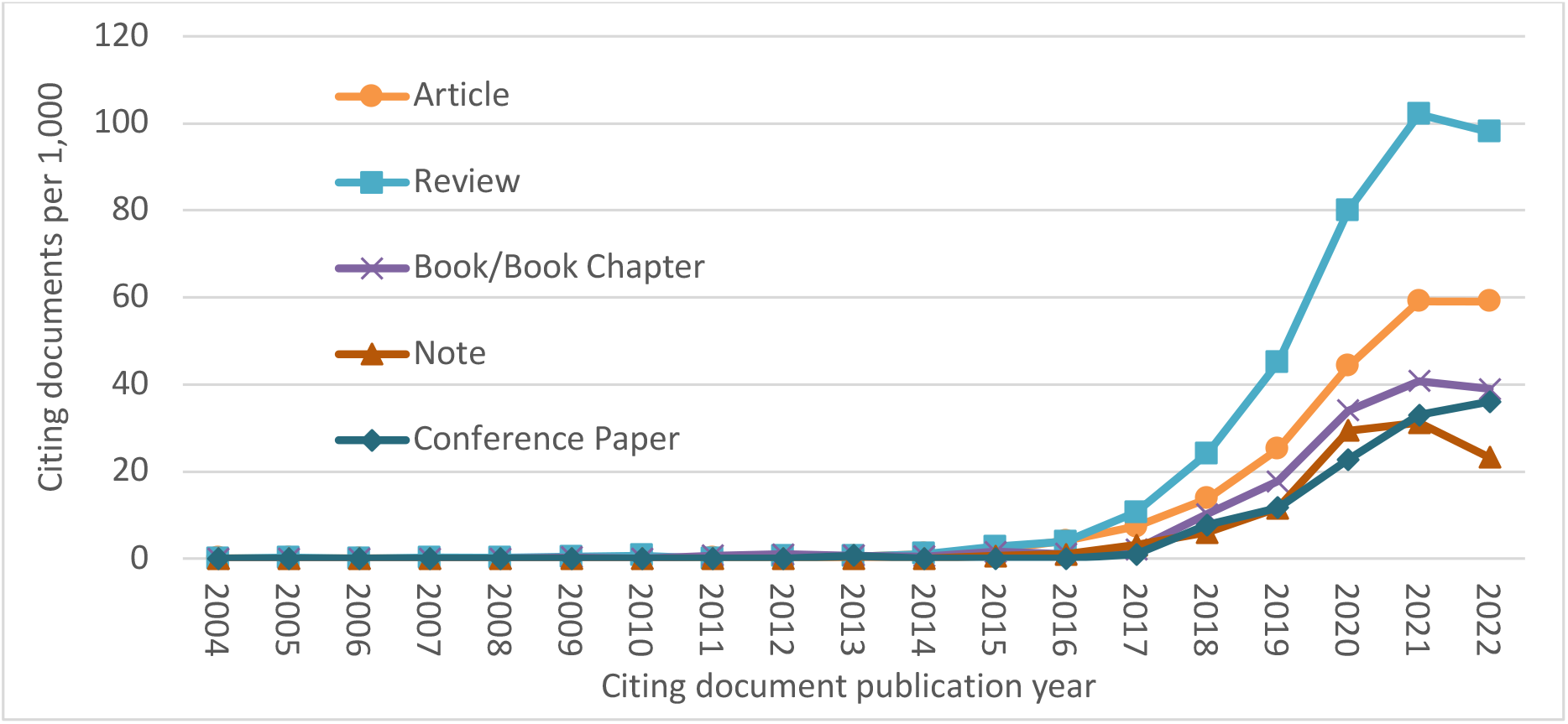
Document types citing preprints per 1,000 Scopus documents.

Figure 2 reveals a dynamic and evolving pattern of citing preprints in different document types within the field of neuroscience from 2016 onward. Notably, reviews seem to be increasingly relying on preprints as references compared to other document types. Journal articles also take the second place in citing preprints. Conference papers, book (chapters), and notes share almost similar rates and show a lower rate of preprint citations.

Figure 3 provides a server-based overview of the number of citations to preprints within Scopus documents in the field of neuroscience.

**Figure 3.**
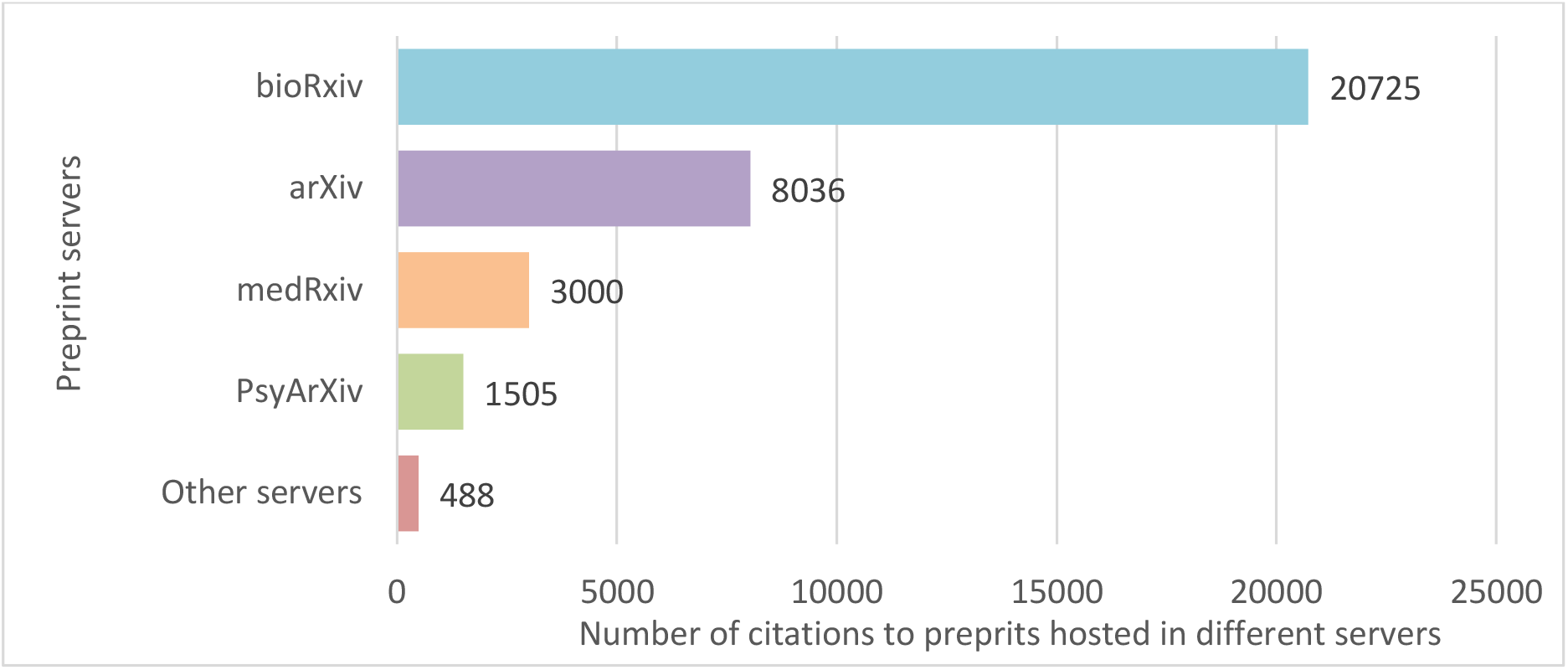
Number of citations to preprint servers within the neuroscience Scopus-indexed documents.

As Figure 3 represents, bioRxiv emerges as the most frequently cited preprint server, with a substantial count of 20,725 citations from all Scopus-indexed neuroscience-related documents. Following bioRxiv, arXiv also commands a significant presence, with 8,036 cited preprints. On the third and fourth places, medRxiv and PsyArXiv have 3,000 and 1,505 cited preprints, respectively. The category “Other servers” encompasses not highly frequently cited preprint servers (i.e., OSF Preprints, SocArXiv, MetaArXiv, EcoEvoRxiv, RePEc, CogPrints, MindRxiv, SportRxiv, ChinArxiv, EdArXiv, Preprints.org, engrXiv, NutriXiv, PaleorXiv, AfricArXiv, BioHackrXiv, EarthArXiv, Frenxiv, INA-Rxiv, LIS Scholarship Archive, and Thesis Commons) with a total of 210 cited preprints.

Figure 4 reflects the trend of citations to preprint servers within Scopus-indexed neuroscience documents. This figure illustrates a better recognition of preprint servers over time, shedding light on neuroscience researchers’ changing attitudes towards preprints.

**Figure 4.**
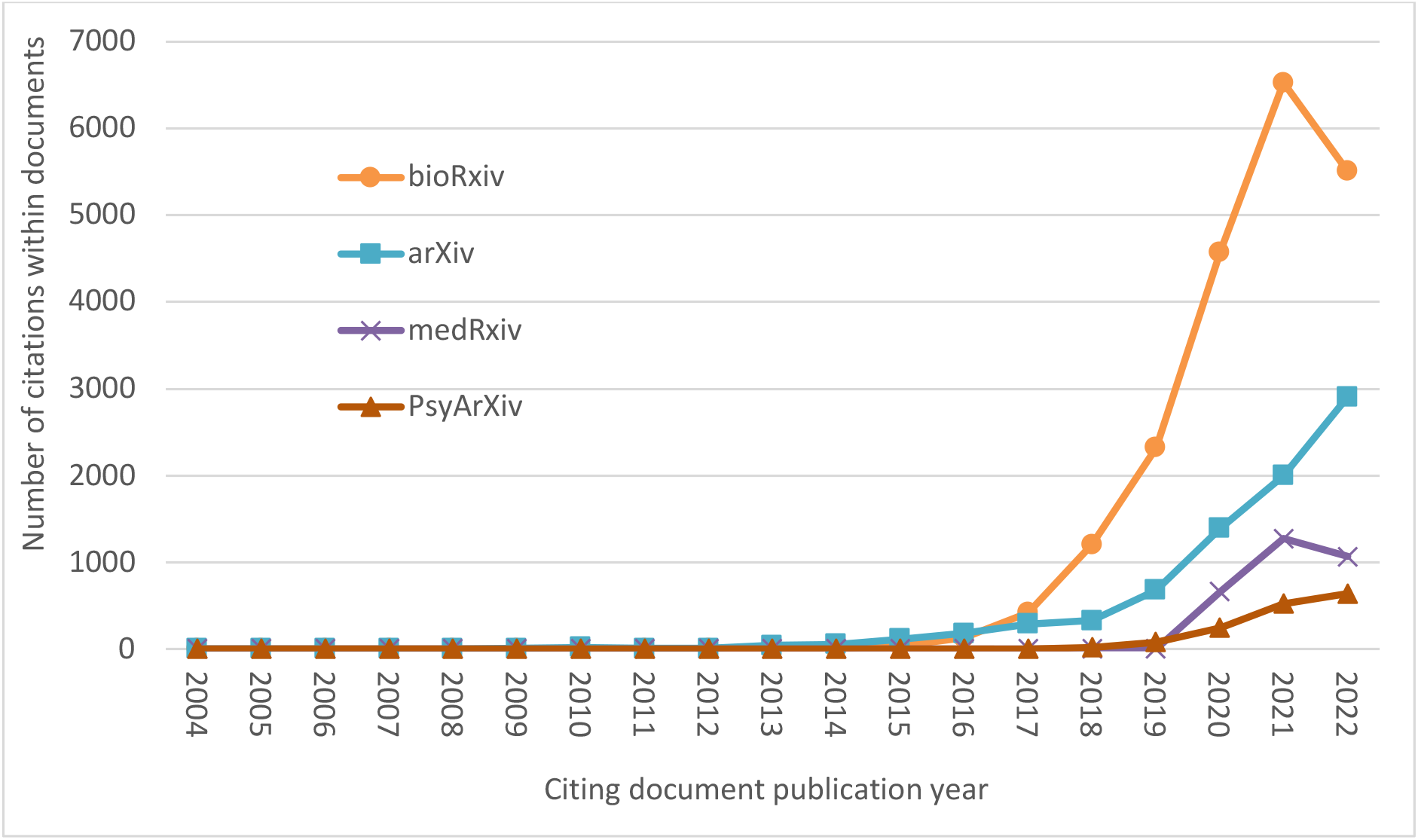
Preprint servers cited within the neuroscience Scopus-indexed documents.

According to Figure 4, regarding the trend of citations to preprint servers, bioRxiv emerged again as being distinctively more frequently cited over years. This easily discernible difference between citations to bioRxiv compared to other servers signifies bioRxiv’s role as an important source for preprints in the field, reflecting the scientific community’s higher confidence in the preprints hosted on this platform. With other servers as well, there is a consistent increase in the number of citations to other preprint servers. This demonstrates that while bioRxiv remains central, neuroscience researchers are also diversifying their input sources, indicating an openness to exploring preprints from different repositories. This broader engagement with various preprint servers also reflects the interdisciplinary nature of neuroscience.

Table 2 provides insights into the top 20 Scopus journals in the field of neuroscience that cite preprints most frequently.

**Table 2.** The 20 Scopus journals citing preprints most frequently.

| Journal | Number of publications citing preprints | Documents citing preprints per 1,000 Journal publications | CiteScore 2022 |
| --- | --- | --- | --- |
| eLife | 1815 | 129 | 12.3 |
| PLOS Computational Biology | 1106 | 116 | 7.1 |
| Frontiers in Neuroscience | 986 | 88 | 6.8 |
| Neuroimage | 654 | 30 | 11.6 |
| Peerj | 539 | 37 | 5.1 |
| Neuron | 532 | 42 | 26.9 |
| Current Biology | 452 | 80 | 12.1 |
| Frontiers in Human Neuroscience | 375 | 46 | 4.4 |
| PLOS Biology | 330 | 54 | 15.4 |
| Frontiers in Neurology | 280 | 23 | 4.8 |
| Human Brain Mapping | 262 | 46 | 9.1 |
| Current Opinion in Neurobiology | 260 | 67 | 11.7 |
| Frontiers in Computational Neuroscience | 254 | 152 | 4.8 |
| Frontiers in Cellular Neuroscience | 236 | 47 | 8.6 |
| Brain Sciences | 232 | 43 | 3.9 |
| Neuroscience and Biobehavioral Reviews | 229 | 41 | 13.4 |
| Journal of Visualized Experiments | 216 | 18 | 2.3 |
| Frontiers in Aging Neuroscience | 211 | 39 | 5.2 |
| Neural Processing Letters | 202 | 87 | 5.4 |
| Trends in Cognitive Sciences | 195 | 67 | 30.4 |

Table 2 presents a journal-based frequency of citations to preprints while facilitating its comparison to the total ratio of documents citing preprints per 1,000 journal publications related to neuroscience. The 2022 CiteScores can have implications of the citability of the journals and their potential roles in the adoption of the preprinting culture.

Table 3 showcases the number of publications within 20 countries with the highest number of publications citing preprints, but also provides context by presenting the ratio of documents citing preprints per 1,000 country publications.

**Table 3.**
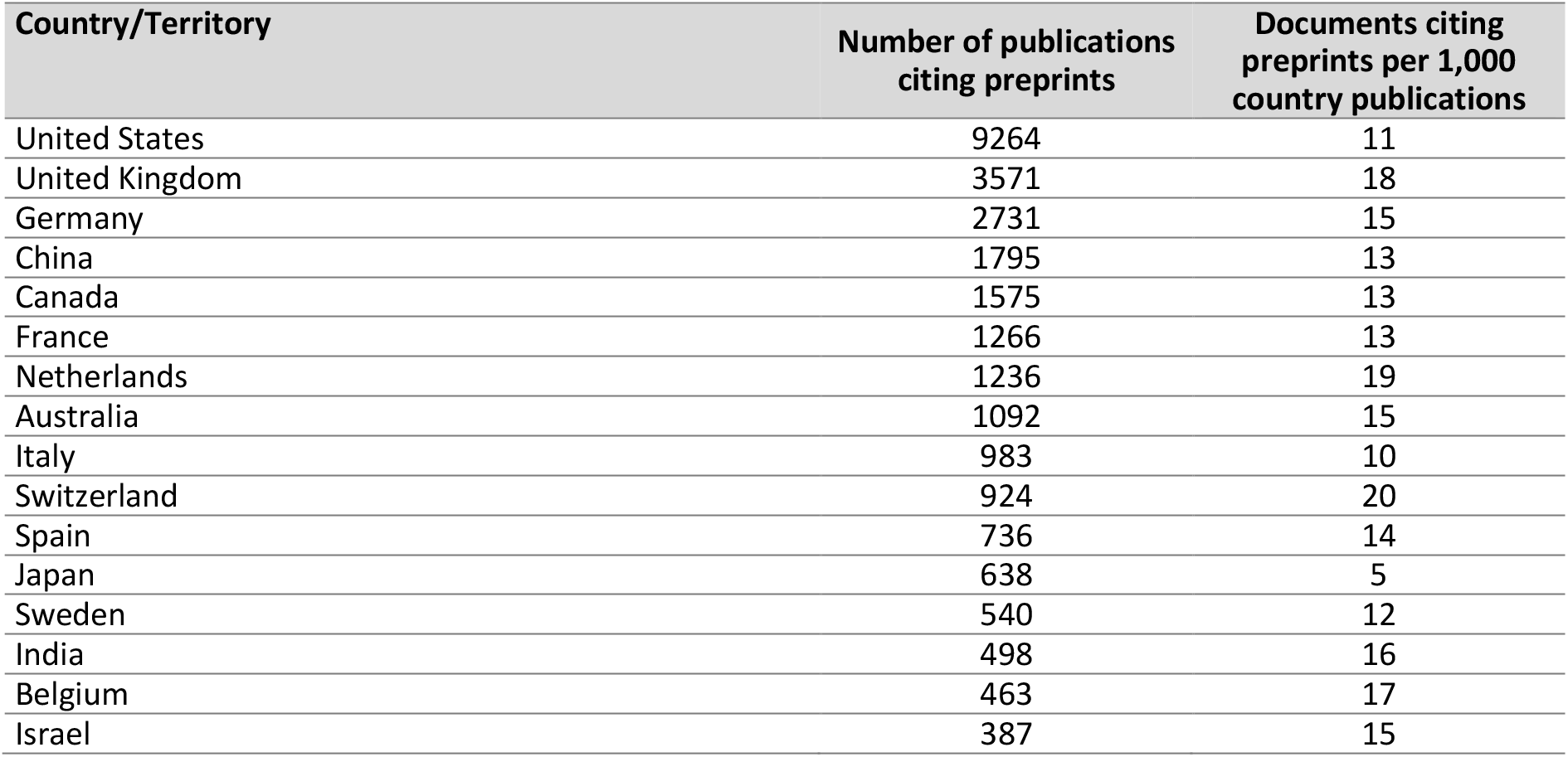
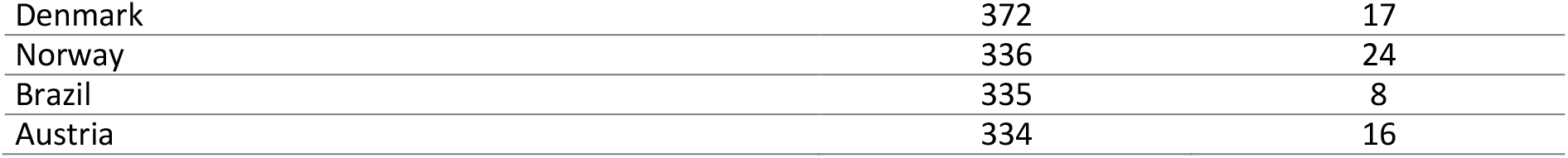
The 20 countries with the highest number of publications citing preprints.

| Country/Territory | Number of publications citing preprints | Documents citing preprints per 1,000 country publications |
| --- | --- | --- |
| United States | 9264 | 11 |
| United Kingdom | 3571 | 18 |
| Germany | 2731 | 15 |
| China | 1795 | 13 |
| Canada | 1575 | 13 |
| France | 1266 | 13 |
| Netherlands | 1236 | 19 |
| Australia | 1092 | 15 |
| Italy | 983 | 10 |
| Switzerland | 924 | 20 |
| Spain | 736 | 14 |
| Japan | 638 | 5 |
| Sweden | 540 | 12 |
| India | 498 | 16 |
| Belgium | 463 | 17 |
| Israel | 387 | 15 |
| Denmark | 372 | 17 |
| Norway | 336 | 24 |
| Brazil | 335 | 8 |
| Austria | 334 | 16 |

While the United States leads the pack with the highest number of publications citing preprints (9,264), a closer look reveals a strong showing from North America with both the US and Canada ranking within the top 5. Additionally, the United Kingdom, Germany, and China are among the top 5. However, when we consider the ratio of documents citing preprints per 1,000 country publications, a distinct trend emerges in Europe. Notably, several European countries, including Norway, the Netherlands, and Switzerland, exhibit comparably higher ratios compared to North America. This suggests a potentially higher level of preprint *adoption* within these European research communities.

To answer our question regarding the motivation behind citations to preprints, Scite identified 1450 citation statements within 451 out of 653 (68% of the research sample) retrievable neuroscience-related publications. Out of the 1450 citations, Scite identified 865 and categorized them as “mentioning”, “supporting”, or “contradicting” (Figure 5). However, the remaining 585 unclassified citations account for 40% of the total mentions which Scite was unable to categorize them.

**Figure 5.**
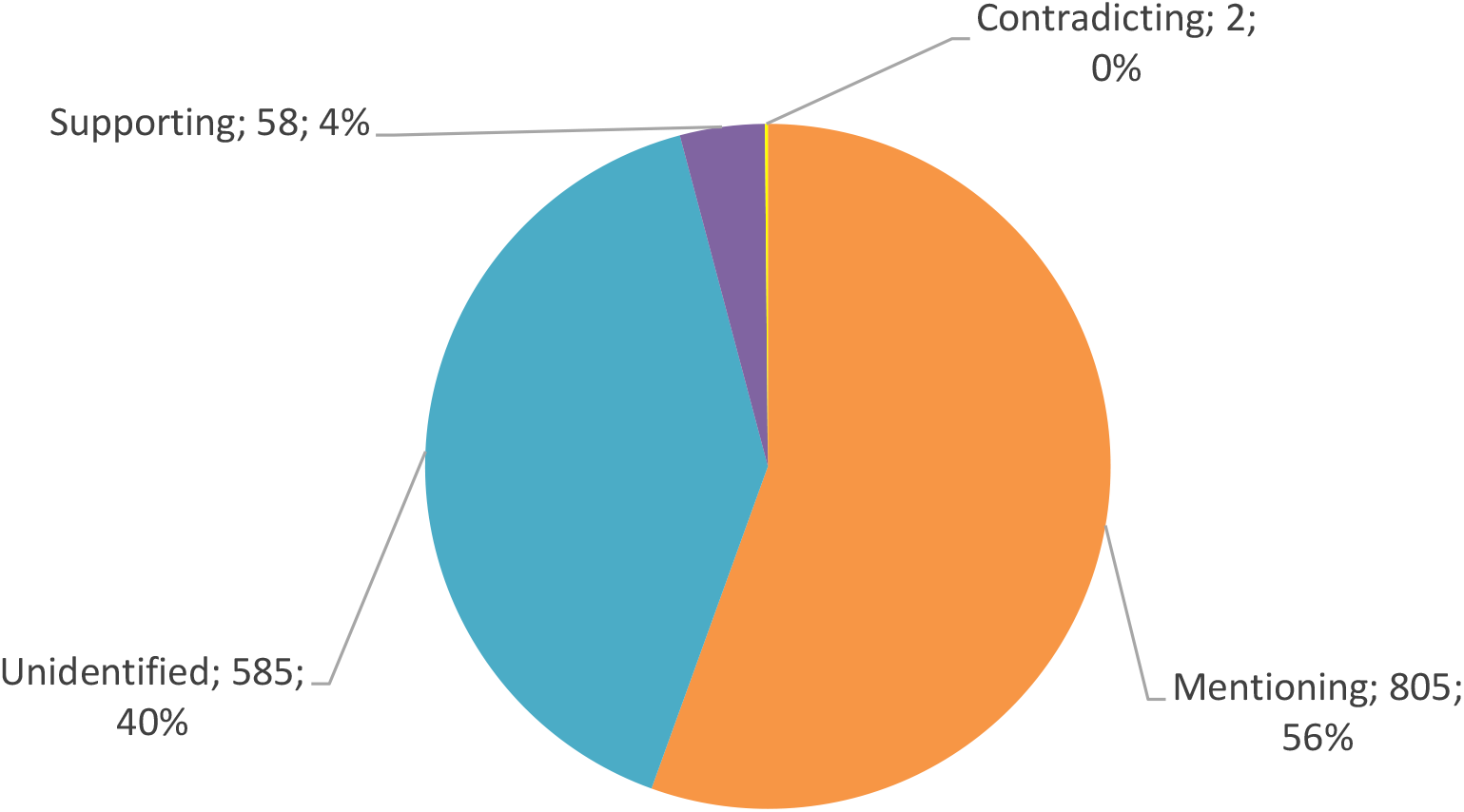
The types of citations to preprints within the neuroscience Scopus-indexed documents.

Figure 5 provides an overview of the nature of preprint citations within a sample of Scopus-indexed neuroscience documents, with “mentioning” being the most prevalent, occurring 805 times. An example of a “mentioning” citation is “To the best of our knowledge, two other methods **have recently been proposed** in [11; a preprint, 12]” (de Weck et al., 2018). However, “supporting” citations are less frequent with 58 instances. An example of a “supporting” citation is “**This is consistent with** conclusions from (Elliott et al., 2019; a preprint) who reported that the test–retest interval had little impact on reliability estimates” (Hassel et al., 2020). Nonetheless, “contradicting” citations are the least common, occurring only two times. An example of a “contradicting” citation is “**In contrast with a recent preprint** (Saldaño et al., 2021; a preprint), the predicted flexibility values failed to correlate with their pLDDT values, …” (del Alamo et al., 2022). As observed in the last example, the lexical content – a recent preprint – can be also informative of the attitude towards preprints.

Regarding the context where preprints are most frequently cited in the Scopus-indexed neuroscience documents, out of the 1450 statement citations found, Scite could only identify the location of 828 within the text of the publications. Data analysis revealed that most publications cited preprints in the “introduction” section (Figure 6).

**Figure 6.**
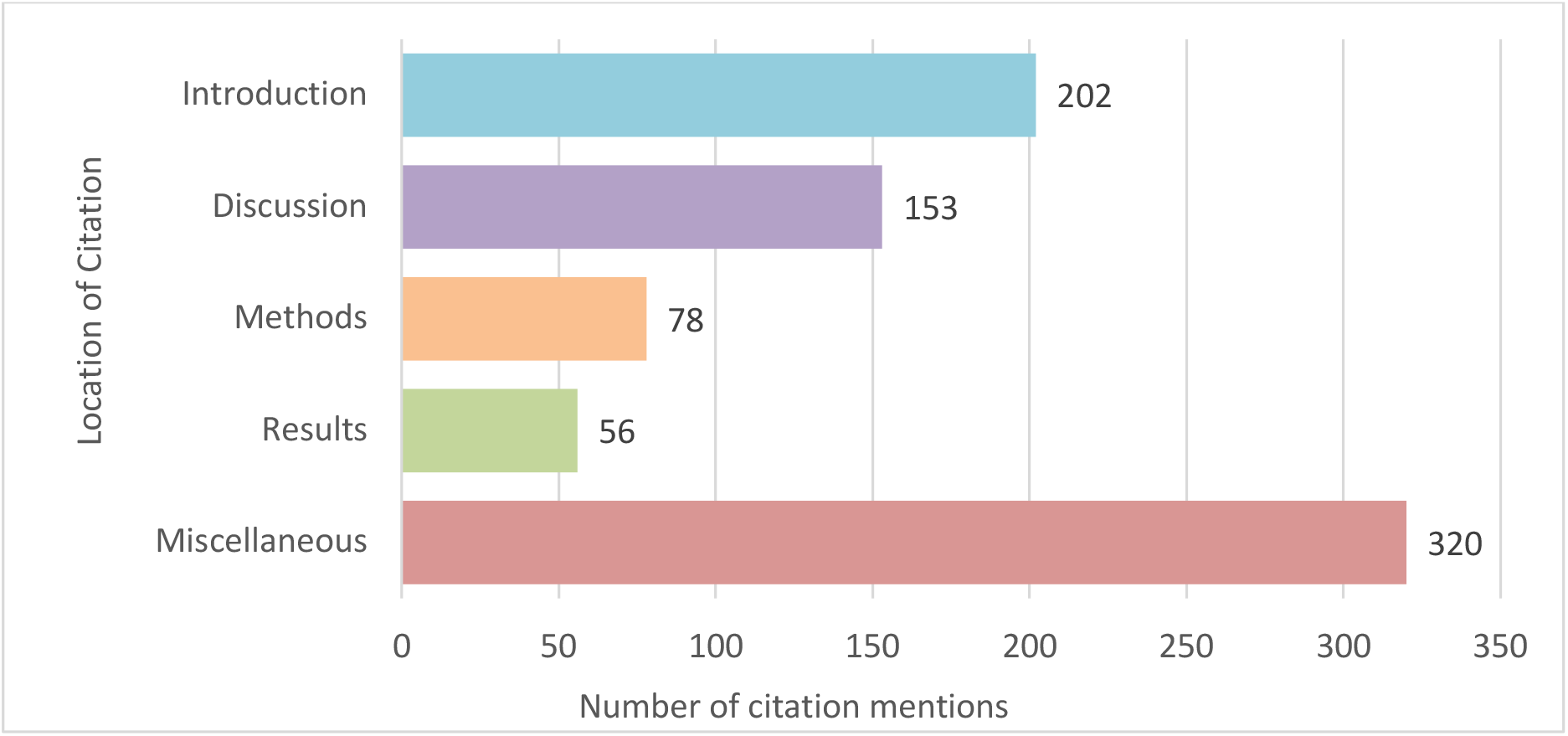
The location of citations to preprints within the Scopus-indexed neuroscience documents.

As seen in Figure 6, most of preprint citations (195; 32.7%) occur in the “introduction” section. Smaller proportions of preprint citations occurred in the “discussion” (153; 25.7%) and the “methods” (78; 13.1%) sections. The location of a considerable number of citations (320; 53.8%), however, is not identified by Scite, so they fall into the “miscellaneous” category.

On a complementary note, the ratio of self-citations to preprints in neuroscience-related Scopus documents were investigated using Scite. Author self-citation refers to citing one’s own previously published work in an upcoming publication (Sri-Ganeshan et al., 2021). Out of 451 records of the research sample found by Scite, 77 (17.07%) were identified as author self-citations to preprints.

## 4 Discussion

The findings of the current study provide a comprehensive insight into the dynamic landscape of preprint citation patterns within the neuroscience domain. These figures unveil the trends of the evolution of the scientific culture among neuroscience researchers.

According to the findings, the early 2000s witnessed a relatively low rate of preprint citations, yet a substantial growth in citing documents became evident from 2014 onwards, peaking in 2021 at around 60 per 1,000 Scopus documents (6%). This trend signifies a shift in how neuroscience researchers engage with preprints. It illustrates a rise in the popularity and recognition of preprints as valid sources of information (Sterling, 2018; Sutton & Gong, 2017), which was accelerated by the spread of Covid-19 leading to a tsunami of preprints (Watson, 2022). This trend is considered to be continuing and changing how science is communicated (Watson, 2022). While until 2014, the realm of preprint servers was confined to a limited number (Li et al., 2015), currently, it has burgeoned into a rich and diverse landscape with numerous servers operating at institutional, national, and global levels. It should be noted though that despite the increase in citations to preprints since their emergence, with only 6% of publications citing preprints in 2021, there is still a long way for preprints to find their way in science communication.

The adoption of preprints in neuroscience can be attributed to the pressing need for up-to-date references in this rapidly evolving field. Neuroscience researchers are often on the cutting edge of knowledge, requiring references that reflect the latest developments and discoveries. Preprints fulfill this demand by offering a channel for researchers to share their findings with the scientific community, making them exceptionally current (Irawan et al., 2022). In contrast, the traditional peer-review process, while valuable for ensuring the quality of publications, can be time-consuming, delaying the dissemination of crucial information. As a result, the immediacy of preprints aligns seamlessly with the dynamism of neuroscience research, enabling researchers to access and build upon the most recent findings in a field where timeliness can be paramount to progress.

One compelling rationale behind the increasing prevalence of preprints in scholarly research is their open accessibility, which addresses multiple critical aspects of academic dissemination. Firstly, preprints are freely accessible, eliminating financial barriers and providing equitable access to knowledge. Researchers, irrespective of their institutional affiliations or financial resources, can readily access and utilize preprints. Furthermore, in scenarios where researchers encounter both a preprint and a restricted access version of a document, the appeal of the open preprint is pronounced. Given the choice between a freely accessible preprint and a version that may require payment or institutional subscriptions, researchers are inclined to opt for the open preprint, ensuring immediate access to the content without any constraints. This not only promotes information equity but also encourages a broader and more inclusive exchange of research findings, aligning with the principles of open science and the democratization of knowledge. This equity is to the benefit of different players in the research domain with early career researchers can take advantage of preprints to be recognized (Rong & Ludo, 2023).

As expected, review articles have cited preprints more frequently compared to other publication types (journal articles, book [chapters], etc.). Considering that a review paper presents the current state of knowledge, the scope of a certain topic, as well as the existing gaps within a specific field in a synthetized and an integrated manner (Palmatier et al., 2018), preprints are inevitably included in a synthesis for a thorough inclusion of the existing knowledge on a certain topic. Second, as preprints enable researchers to immediately disseminate their new discoveries and innovative findings, they are included in the review papers as unique insights and novel methodological approaches are supposed to be evaluated in review papers (Palmatier et al., 2018). Besides, review papers are at times allowed to have more references, so this might be another reason why there is a higher probability of citing preprints in reviews. On a positive note, review papers discuss inconsistent findings in the literature (Palmatier et al., 2018), so with the “unreliability misconception” in relation to preprints, this can be to the benefit of the preprinting culture.

Journal articles also have been citing preprints more frequently and consistently during the past decade. Considering that authors of a publication could be reviewers for another piece, citation to preprints could imply a careful analysis and examination of the cited preprint. Moreover, a peer-review has further analyzed the content of the publication citing a preprint, which could be considered as a double investigation of inconsistencies.

On the other end of the spectrum, books (chapters) remain notably hesitant to incorporate preprints as references, pointing to a marked difference in their adoption compared to other document types. This divergence may be attributed to the inherent goals and content of this category. Book (chapters) typically do not hinge on the urgency of referencing the latest research findings. Instead, they often prioritize citing sources with a well-established and accredited reputation, given their inclination towards comprehensive and enduring knowledge. This selectivity in the reference selection reflects a level of caution among book (chapter) authors, as preprints, lack the traditional peer-review. Consequently, book (chapter) authors may exhibit reservations about relying on the content of preprints, preferring the reliability and credibility offered by peer-reviewed and well-established sources. This marked distinction underscores the varied information needs and citation practices within the academic landscape, acknowledging the enduring role of traditional publishing in certain scholarly contexts. From another perspective, sometimes it takes years to publish a book, thus we speculate that with the proportional rise in citations to preprints, more preprints will be cited by book (chapter) authors.

When it comes to preprint citations, neuroscientists distinctly prefer bioRxiv, ArXiv, medRxiv, and PsyArXiv servers. This preference is intricately tied to the interdisciplinary nature of neuroscience, a field that seamlessly bridges the realms of medicine, biology, psychology, and technology. Moreover, based on the number of records registered on each preprint server (see Table 1), it seems some preprint servers are more trusted or even better known and considered to be of higher quality.

An examination of the journals frequently citing preprints reveals a particularly striking trend – authors publishing in eLife exhibit a substantial predilection for preprints. This phenomenon can be attributed to a notable policy outlined in the journal’s author guide stipulating that “*eLife only peer-reviews submissions that are available as preprints*” and “*reviewers’ reports would be published alongside the paper, together with a short editorial assessment of the work’s significance and rigour*” to combine “*the immediacy and openness of preprints with the scrutiny of peer-review by experts*” (https://web.archive.org/web/20231006105925/https://elife-rp.msubmit.net/html/elife-rp_author_instructions.html#process). This unique requirement has evidently a profound influence on authors’ research behavior, as it not only signifies a journal’s strong endorsement of preprints but also encourages authors to actively engage with preprints. In essence, eLife’s policy serves as a representative example of how a journal’s policy can significantly shape research practices and foster a culture of preprint adoption. On the other hand, the SLEEP® journal has clearly pointed out that it “*does not allow citation of preprint manuscripts in final published articles. Prior to publication of accepted papers, preprint citations must be replaced with the final, peer-reviewed version of record. If the cited preprint work has not been published by acceptance, it must be removed from the reference list*.” (https://academic.oup.com/sleep/pages/General_Instructions). As mentioned above, while the changes to a preprint after it is published as a peer-reviewed publication are a legitimate concern, the findings of a recent study on Covid-19-related RCTs has shown that the main results and conclusions remain the same in 98% of peer-reviewed preprints (Bai et al., 2023). Along the same lines, another study on the bioRxiv and medRxiv showed changes in the conclusions of 17.2% of Covid-19-related and 7.2% of non-Covid-19-related preprints after being peer-reviewed, yet most of the changes did not induce significant qualitative differences between the two versions (Brierley et al., 2022). The implicit message here is that for a wider promotion of preprints, journals should adopt more proactive policies to encourage authors to embrace preprints as a vital element of scholarly communication and thus contribute to the broader open science movement. In fact, the open peer-review process in some journals (e.g., eLife) is playing an important role in promoting transparency, yet the high article processing charges still function as barriers. Instead, public commenting on preprints while citing a preprint could be very helpful in the science communication on a broader level.

Furthermore, the inclusion of journals 2022 CiteScore reveals the influence of highly credible journals. “Neuron”, for example, emerges as a standout with an exceptional CiteScore of 26.9 and among journals with the highest number of publications citing preprints, underscoring its profound impact on the neuroscience community. The presence of several other high-impact journals in this list, like “PLOS Biology” and “Trends in Cognitive Sciences”, reaffirms the importance of preprints in disseminating scientific findings to a wider audience.

Among the 20 countries with the highest number of publications citing preprints, the regional patterns of citations to preprints represented North America, China, and Europe as prominent users. This preference can be attributed to a multifaceted interplay of factors, including the influential role of funding policies that formally recognize and endorse preprints in these regions, as evidenced by studies like Kaiser (2017). Notably, the European continent has exhibited a significant commitment to open science, with the European Commission recently spearheading the development of policies supporting open research practices (European Commission, 2023). On the other hand, policies against preprints; for example, the recent changes regarding the preprint ban in grant applications initiated by the Australian Research Council (ARC) (Ciriminna et al., 2021), can decrease preprint utilization by researchers.

In terms of the motivations behind citations to preprints, “mentioning” citations, by far the most frequent, showcase preprints as essential sources of neuroscience literature, contributing to identifying research gaps and laying the foundations for future research endeavors. Besides, “supporting” citations, although less frequent, highlight the endorsement of preprint findings, emphasizing their increasingly important role in shaping the scientific discourse. Interestingly, “contradicting” citations, with the lowest frequency, suggest increasing levels of trust in preprints and shows they are recognized as sources of information to be discussed.

According to the findings, it is evident that neuroscience researchers hold preprints in high regard to a greater extent recently, seamlessly incorporating them into their scholarly discourse alongside other forms of academic literature, including book (chapters), conference papers, and peer-reviewed journal articles. In addition, the current study revealed a prevalence of neuroscience researchers citing preprints throughout their publications, with a particular emphasis in the introduction section. It is important to note that introductions typically contain more citations than other sections to establish context and prior research (Bertin & Atanassova, 2022). However, Bertin and Atanassova’s (2022) research on examination of PLOS journal citations indicated a higher frequency of preprint citations within the methodology section. Note that the current research considers absolute number, but Bertin and Atanassova’s (2022) research considered relative number (i.e. the number of citations to preprints within the methodology section divided by the total number of citations within the methodology section).

Regarding self-citations, our findings indicated that 17% of citations to preprints were self-citations. A previous study has reported an overall self-citation rate of 9% across disciplines (varying from 3% to 15%) (Szomszor et al., 2020). Along the same lines, the ratio of neuroscience researchers’ self-citations to preprints is comparable to that of physical sciences (15%). Considering that funding/research organizations and universities require open access publication of study protocols and findings and regarding researchers’ willingness to show productivity and to be recognized, authors might opt for preprints to fulfill these needs. This finding needs further investigation in future research to understand the reasons behind the higher prevalence of self-citation within the context of preprint.

The findings of our study are neither temporally stable nor generalizable across fields as the Covid-19 pandemic significantly boosted the production of scientific publications and preprints, initially leading to a 60% surge in submissions to Elsevier journals and a tenfold increase in submissions on medRxiv (Jocalyn, 2023). For instance, recent studies are being published on neurological complications of Covid-19 (Needham et al., 2020; Spatola et al., 2022), therefore this is probably exclusive to the contemporary era and to the health care field. Furthermore, considering that massive amounts of data have been manipulated and processed in this study using python codes (Rasuli et al., 2023), some citations might have been missed in our final dataset. Some journals (such as Sleep) do not allow referencing to preprints; thus, they are automatically excluded from the current study. Finally, in line with the heated discussions on the contributions and perils of artificial intelligence-based tools, Scite significantly accelerated the data analysis procedure although it could not be fully automatized and needed further manipulations; however, there were a lot of missing information as the context of the citations were not correctly identified for a large number of records. These tools can be, thus, helpful in automating tasks, increasing efficiency, and saving time, yet still need to be significantly improved.

Similar studies can broaden our understanding of the current and future contributions of preprints to scientific research. First and foremost, a deeper analysis of self-citation patterns in citing preprints could be highly informative of what objectives authors are pursuing by citing preprints. For instance, many senior and junior researchers might publish preprints to be able to cite them in their grant applications but not to provide the community with unconstrained accessibility. Furthermore, since different lexical contents are used for citing preprints (Bertin & Atanassova, 2022), linguistic content analysis of these lexical contents can be informative of both the trust put in preprints and the credibility of preprints in the scientific community. Moreover, since early career researchers can publish preprints to show productivity and receive recognition (Rong & Ludo, 2023), a closer look at the scientific profiles of the authors of preprints can shed light on the contributions of preprints as who they cater for the most in the scientific community. Additionally, in a recent survey on familiarity with preprints (Rong & Ludo, 2023), it was highlighted that 55-96% of researchers (in disciplines except mathematics, computer science, physics and astronomy) never read a preprint. Thus, a survey on attitudes towards publishing and/or citing preprints can be interesting as reasons behind not adopting the preprinting culture might not be that they are unreliable, but that they are not well-recognized or at least were not to a majority before Covid-19.

As a conclusion, the findings suggest that preprints play a multifaceted role in the research process, catering to neuroscience researchers at various stages of their research. Along with other sources of information, they serve as valuable resources for those seeking to contextualize and formulate research ideas. Preprints are equally instrumental in performing a literature review, helping neuroscience researchers keep abreast of the latest developments in the field with the unstoppably fast advances.

## 5 Acknowledgements

The study was supported by the University and University Hospital of Liège, the Belgian National Funds for Scientific Research (FRS-FNRS), the FNRS MIS project (F.4521.23), FNRS PDR project (T.0134.21), the ERA-Net FLAG-ERA JTC2021 project ModelDXConsciousness (Human Brain Project Partnering Project), the fund Generet, the King Baudouin Foundation, the BIAL Foundation, the Mind Science Foundation, the European Commission, the Fondation Leon Fredericq, and the Horizon 2020 MSCA – Research and Innovation Staff Exchange DoC-Box project (HORIZON-MSCA-2022-SE-01-01; 101131344).

